# Large Language Models Reveal the Neural Tracking of Linguistic Context in Attended and Unattended Multi-Talker Speech

**DOI:** 10.1101/2025.04.24.648897

**Authors:** Corentin Puffay, Gavin Mischler, Vishal Choudhari, Jonas Vanthornhout, Stephan Bickel, Ashesh D. Mehta, Catherine Schevon, Guy M. McKhann, Hugo Van hamme, Tom Francart, Nima Mesgarani

**Affiliations:** KU Leuven, Dept. Neurosciences, ExpORL, Leuven, Belgium; KU Leuven, Dept. of Electrical Engineering (ESAT), PSI, Leuven, Belgium; Columbia University, Dept. of Electrical Engineering, New York, NY, USA; Hofstra Northwell School of Medicine, Uniondale, NY, USA; The Feinstein Institutes for Medical Research, Manhasset, NY, USA; Columbia University, Dept. of Neurology, New York, NY, USA; Columbia University, Dept. of Neurological Surgery, Vagelos College of Physicians and Surgeons, New York, NY, USA

**Keywords:** Auditory attention, Large Language Models, Context Encoding

## Abstract

Large language models (LLMs) capture long-range contextual structure in natural language and have recently been shown to align with the human brain’s contextualized linguistic encoding. This makes them a promising computational probe for studying how context-dependent linguistic information is represented during natural speech perception. Speech perception often occurs in multi-talker environments, where attention must dynamically select among competing streams, yet how contextual information from attended and unattended speech is neurally encoded remains underexplored. Here, we investigate how auditory attention modulates neural tracking of context-dependent linguistic representations using electrocorticography (ECoG) and stereoelectroencephalography (sEEG) recordings from three epilepsy patients engaged in a two-conversation “cocktail party” paradigm. To model neural responses to attended and unattended speech streams, we used contextual word embeddings generated by large language models. We find that LLM-derived features reliably predict brain activity for the attended stream and that contextual information from the unattended stream also contributes to neural prediction. Importantly, these contributions extend beyond low-level acoustic features and shallow syntactic information, and depend on the surrounding linguistic context. Moreover, neural tracking of the unattended stream reflects shorter-range contextual integration than that of the attended stream. Together, these findings indicate that neural responses to speech reflect context-dependent linguistic representations from multiple concurrent speech streams, with attention modulating the depth and timescale of contextual integration. Our results highlight the utility of LLMs for probing higher-level linguistic representations in complex, naturalistic listening environments.

## 1 Introduction

The emergence of Large Language Models (LLMs) and their high performance on natural language processing tasks has attracted increasing interest from neuroscience. LLMs capture long-range contextual structure in language, and recent work has shown that their representations align with human brain activity during language processing (Caucheteux et al., 2023; Goldstein et al., 2022; Mischler et al., 2024; Schrimpf et al., 2021). This has motivated the use of LLM-derived embeddings as computational probes to study how language is encoded in the brain. A common approach is to extract word representations from LLMs and train linear regression models to predict corresponding neural responses (Abnar et al., 2019; Anderson et al., 2021; Caucheteux & King, 2022; Hosseini et al., 2024; Schrimpf et al., 2021; Sun et al., 2021; Toneva & Wehbe, 2019). The correlation between predicted and observed neural activity serves as a measure of how well the model’s representations align with brain responses and allows researchers to investigate which aspects of language (i.e., lexical, syntactic or semantic) are reflected in the brain response. However, LLM-derived embeddings contain multiple types of information: intrinsic properties of the word itself (e.g., lexical identity, frequency) as well as contextual information from preceding words. To disentangle these contributions, some studies have analyzed LLM activations using controlled manipulations of naturalistic transcripts (Caucheteux et al., 2021), while others have compared contextual embeddings to non-contextual embeddings derived from static models like GloVe (Goldstein et al., 2022; Pennington et al., 2014).

Most prior studies have examined neural encoding of language in controlled, single-speaker paradigms. Real-world listening, however, frequently involves multiple talkers, requiring attention to dynamically select one speech stream over others. Multi-talker environments introduce additional challenges for neural encoding: while low-level acoustic features are robustly tracked even under noisy conditions (Ding & Simon, 2012; Lalor & Foxe, 2010), higher-level linguistic representations rely on abstract, spatially localized, and temporally integrated processes (Hickok & Poeppel, 2007; Poeppel et al., 2009). Reduced acoustic signal-to-noise ratio can disproportionately disrupt these higher-level representations (Peelle et al., 2010). Noninvasive EEG studies suggest that attention is necessary for encoding contextual language features, with unattended speech typically showing minimal predictive power (Anderson et al., 2024; Brodbeck et al., 2018; Broderick et al., 2018). However, the low signal-to-noise ratio of EEG limits our ability to determine whether unattended speech is truly not encoded or simply not detectable. Invasive recordings such as electrocorticography (ECoG) and stereoelectroencephalography (sEEG) offer higher fidelity and spatial precision, enabling direct measurement of local high-frequency activity and more accurate characterization of context-dependent linguistic processing in complex listening scenarios (Chang et al., 2010; Mesgarani et al., 2014). These techniques allow us to investigate how attention shapes the encoding of contextual information in both attended and unattended speech streams.

Here, we use LLM embeddings to model neural responses measured with ECoG and sEEG while participants listen to two concurrent speech streams. First, we compare prediction scores across LLM layers for attended and unattended speech, showing that both streams predict the brain response, with higher power for the attended stream. Second, we manipulate the contextual input provided to the LLM, demonstrating that deeper layers improve prediction only when the preceding context matches the actual speech. Third, by varying the length of the context fed to the LLM, we show that neural responses track shorter-range context for unattended speech than for attended speech. Finally, we show that LLM embeddings predict neural activity beyond low-level acoustic features. Together, these analyses reveal how attention shapes the depth and timescale of contextual linguistic encoding in the human brain.

## 2 Results

To explore how linguistic context is represented in the human brain during naturalistic listening and selective attention, we analyzed intracranial recordings from three epilepsy patients as they performed a dual-speaker auditory attention task (Figure 1). Participants listened to two overlapping conversations and were instructed to attend to one, while their neural responses were recorded via sEEG and ECoG electrodes implanted over the left hemisphere of their brain. High-gamma (70 - 150 Hz) neural activity was extracted and used as a proxy for local neural engagement (Edwards et al., 2009).

**Figure 1:**
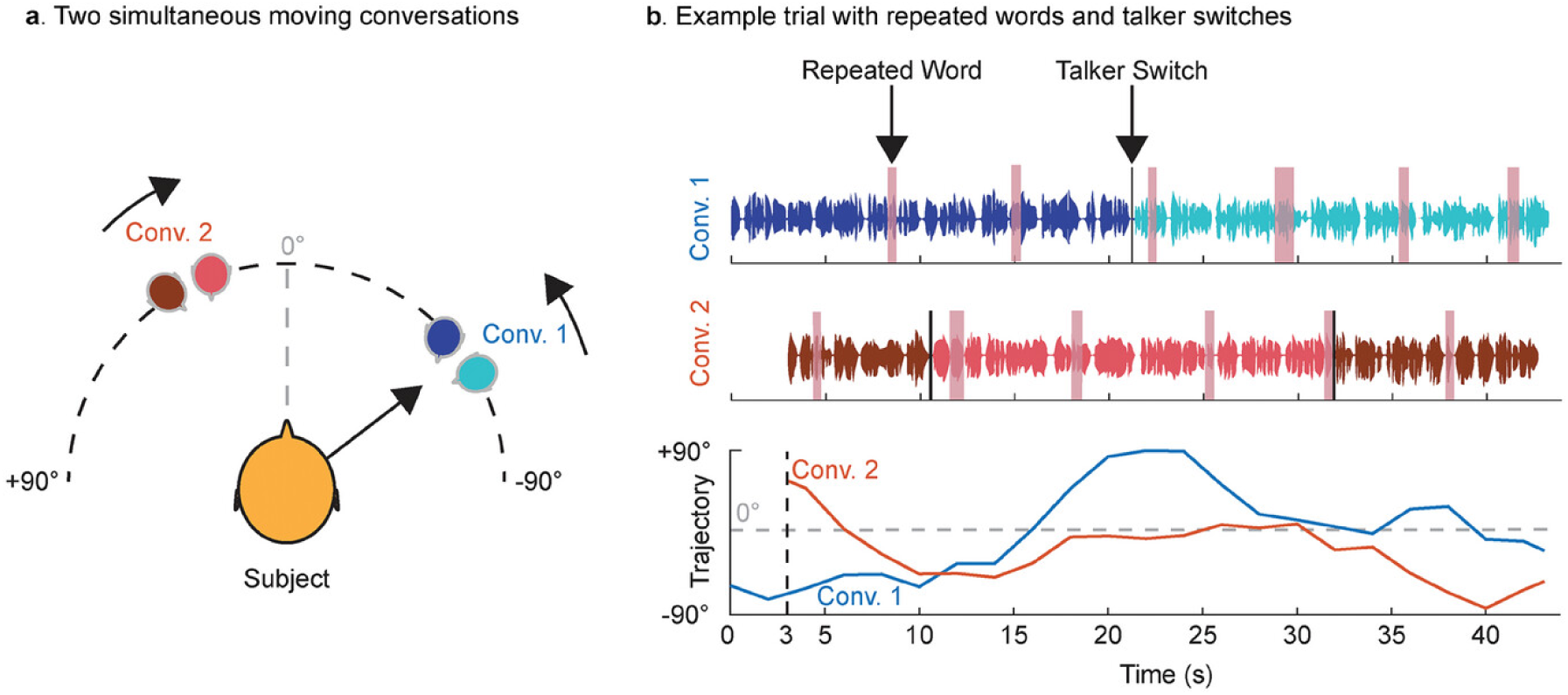
Auditory attention experiment design from Choudhari et al., 2024. (a) Each trial consisted of two concurrent conversations presented simultaneously in the frontal hemifield. The two conversations (Conv. 1 and Conv. 2) moved independently and each involved two distinct talkers who alternated speaking. (b) In each conversation, talkers occasionally repeated a word at random intervals during a 1-back detection task (highlighted in pink). In the cued (attended) conversation, a single talker switch occurred at approximately 50% of the trial duration, whereas the uncued (unattended) conversation contained two talker switches, occurring at approximately 25% and 75% of the trial duration.

To model the contextual encoding across attentional states, we used the Mistral-7B LLM to generate word representations from attended and unattended speech transcripts (Figure 2). Using ridge regression, we quantified how well these embeddings predicted neural responses across different model layers and temporal lags, as well as across varied contextual input given to the LLM. This framework enabled us to examine how attention, contextual input, and context length shape the brain’s alignment with LLM-derived representations.

**Figure 2:**
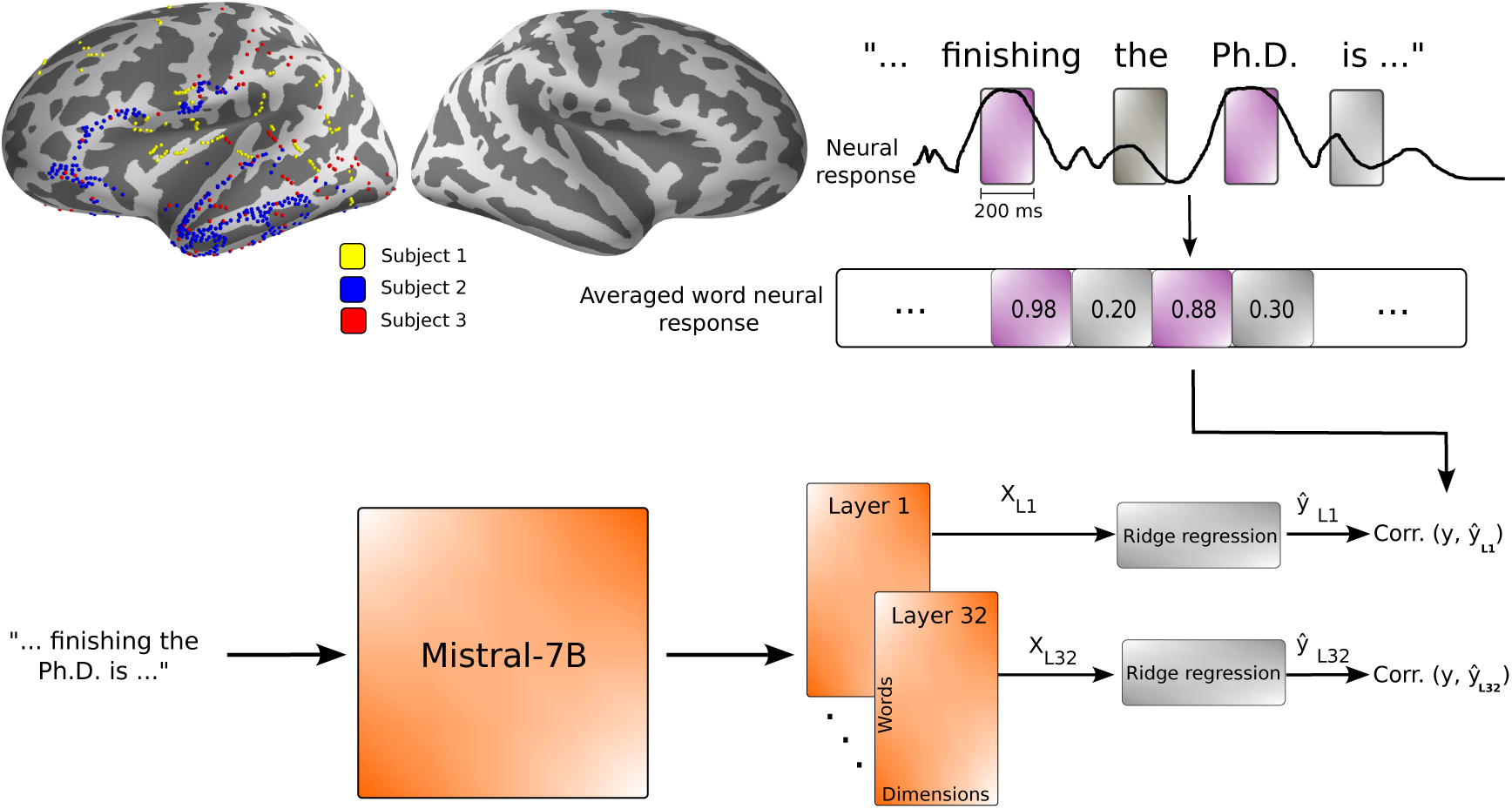
LLM embeddings to predict brain responses. (Inspired from Mischler et al., 2024). We averaged the neural responses over 200-ms windows centered at seven lags, from 0 to 500 ms after word onsets. The word representations are extracted using different layers of Mistral-7B to further predict the averaged word response. The correlation between the predicted and actual responses is calculated to compute the brain score.

In what follows, we present four key findings: (1) attention enhances brain prediction scores from LLM embeddings, (2) this effect is primarily driven by contextual information, (3) the encoded context length is shorter for the unattended than the attended speech, and (4) the information contained in LLM embeddings and predictive of the brain response is not merely acoustic but rather reflect contextual semantic information. Together, these results highlight attentional modulation of linguistic context processing in the brain.

### 2.1 Result 1: Language encoding, measured with LLM brain prediction scores is impacted by attention

In Figure 3, we depict the LLM word representation prediction scores for every layer for both the attended and unattended speech transcripts. The attended and unattended prediction scores for subjects 1, 2, and 3 are depicted in Figures 3a, 3c, and Figure 3e, respectively. We observe a significant difference in the averaged brain score magnitudes across layers between the attended and unattended conditions (two-sided Wilcoxon signed-rank test, *W* = 1626 − 1996, *p* < 0.001, Bonferroni-corrected *α* = 0.05/6, for subjects 1, 2 and 3, respectively), suggesting differences in the language information encoded across attention states. The median difference was 0.023, 0.047, and 0.010 for subjects 1, 2 and 3, respectively, consistent with a higher prediction score for the attended condition.

**Figure 3:**
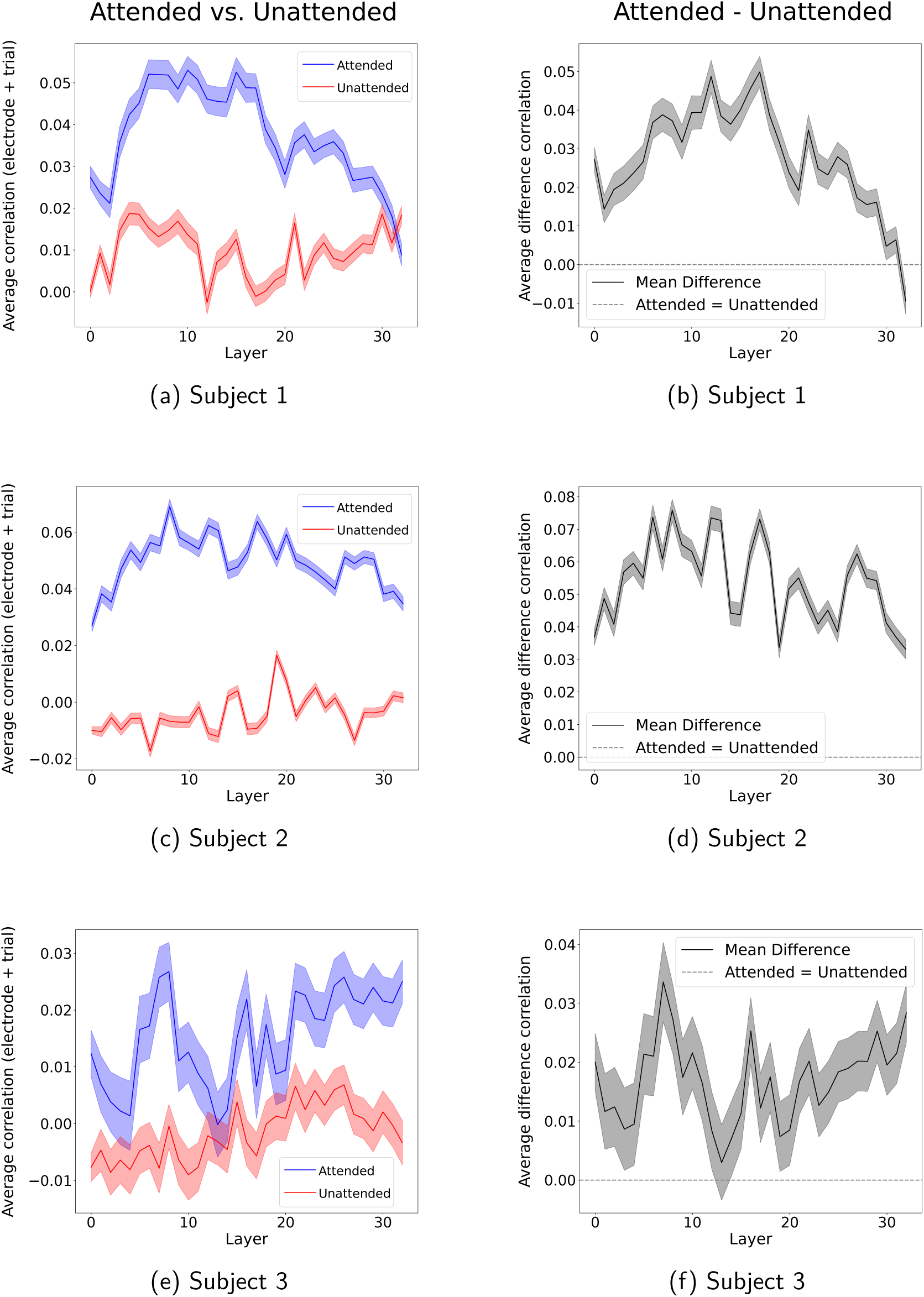
Brain prediction scores of LLM word representations across layers with attention modulation. In (a), (c), and (e) (for subjects 1, 2, and 3, respectively), the two curves depict the average brain score across trials and electrodes for each layer, with SEM across trial-averaged electrodes. Blue: attended speech transcript; red: unattended. (b), (d), and (f) show the attended-unattended score differences.

By definition, the output of Mistral-7B’s layer 0 corresponds to token and positional embeddings only, without any contextual integration, while higher layers progressively integrate greater contextual information (Voita et al., 2019). Thus, as suggested in (Caucheteux et al., 2021), improvements in brain reconstruction scores observed in subsequent layers likely reflect the accumulation of contextual information. We observe that the attended condition shows a difference in prediction score, e.g., between layer 0 and layer 6 (two-sided Wilcoxon signed-rank test, *W* = 1385, 4253, *p* < 0.001, Bonferroni-corrected *α* = 0.05/6) for subjects 1 and 2. For all subjects, a prediction score difference was observed between layer 0 and 7 (two-sided Wilcoxon signed-rank test, *W* = 1380 − 4123, *p* < 0.001, Bonferroni-corrected *α* = 0.05/6) and between layer 0 and 8 (two-sided Wilcoxon signed-rank test, *W* = 1336 − 3246, *p* < 0.001, Bonferroni-corrected *α* = 0.05/6). For layer 6, subjects 1 and 2 showed median differences of 0.023 and 0.028, respectively. Additionally, the median difference subject ranges were 0.015-0.026 and 0.012-0.044 for layers 7 and 8, respectively. These values are consistent with a higher prediction score for these layers compared to layer 0. Results were similar for neighboring layers. This indicates that the contextual information contained in word representations from layers 6, 7, and 8 for subjects 1, 2, and 3, respectively, enables a better prediction of the average brain response than layer 0.

In comparison, we also observed a difference in prediction score for the unattended transcript in subject 1 between layer 0 and 6 (two-sided Wilcoxon signed-rank test, *W* = 2917, *p* < 0.001, Bonferroni-corrected *α* = 0.05/6). In addition, the median difference was 0.015, consistent with a higher prediction score for layer 6 compared to layer 0, and results were similar for neighboring layers. On the other hand, this effect was not found for subjects 2 and 3. While a difference was observed for the unattended case in subject 2 from layer 0 to 6 (two-sided Wilcoxon signed-rank test, *W* = 21951, *p* = 0.24), the median difference was −0.01, consistent with a lower prediction score for layer 6 compared to layer 0. Neighboring layers did not show any difference in prediction scores with layer 0. Subject 3 did not show any significant prediction score between layer 0 and layers 6, 7, 8 (two-sided Wilcoxon signed-rank test, W=2606-3176, p=0.04-0.65, not significant after Bonferroni correction, *α* = 0.05/6). This finding suggests that there might be an unattended context encoding for subject 1.

Figures 3b,3d, and 3f depict the attentional modulation of electrodes (i.e., the brain prediction score of the attended condition minus that of the unattended condition).

We did not observe any localized effect of attention when looking at the prediction scores across brain locations. To support this statement, we provide brain mappings of correlations across subjects when looking at the benefit of attention, i.e., the difference in prediction scores between the attended and unattended conditions at layer 10 (see Supplementary Figure 1).

In the Supplementary Material, we run three additional analyses: (1) to evaluate the impact of repeated words on the prediction score (see Text S1, and Supplementary Figure 2), (2) to evaluate any speaker-specific effects that our model could learn and that could impair our conclusions (see Text S2, and Supplementary Figure 3), and (3) to evaluate the impact of attention drops measured behaviorally (see Text S3). None of these potential confounds changed the results of our analyses.

### 2.2 Result 2: Improvement of LLM prediction scores across layers reflects contextual encoding

To verify that the prediction score’ increase across Mistral’s layers is due to contextual information, we investigate how modifying the context input provided to Mistral influences its ability to predict brain responses. Thus, we repeat the analysis of the previous subsection. However, this time, we either give Mistral the original trial context before the current token (i.e., the one presented to the participants), as done in the previous subsection, or a completely unrelated context extracted from a different story but of the same length, so that it would not be helpful for the LLM to predict the next word (see Context modulation Section for details). We depict the resulting prediction scores for the original and unrelated context conditions in both the attended and unattended conditions in Figures 4a, 4c, and 4e for subjects 1, 2, and 3, respectively.

**Figure 4:**
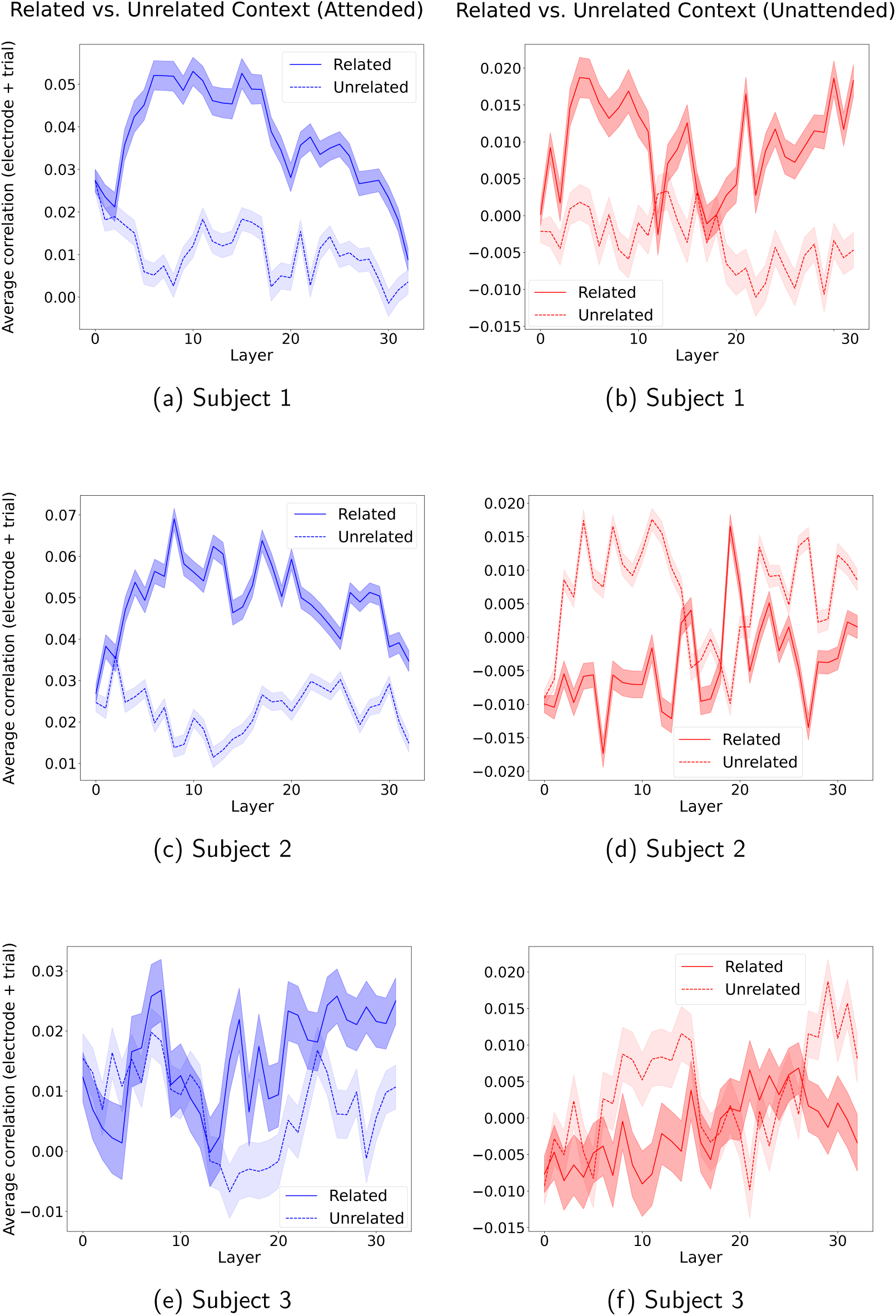
Related vs. unrelated context prediction scores across Mistral layers for both attended and unattended conditions. (a), (c) and (e) depict (for subjects 1, 2, and 3, respectively) the related context (blue curve) and unrelated context (light blue curve) trial-electrode averaged prediction scores across Mistral layers when using the attended word representations. (b), (d) and (f) depict (for subjects 1, 2, and 3, respectively) the related context (red curve) and unrelated context (light red curve) trial-electrode averaged prediction scores across Mistral layers when using the unattended word representations.

In the attended case, we observed a difference in the averaged prediction scores across layers between the related and the unrelated conditions for all subjects (two-sided Wilcoxon signed-rank test: *W* = (562 - 1999, *p* < 0.001, Bonferroni-corrected *α* = 0.05/6, for subject 1, 2 and 3, respectively). The difference of medians between the related and the unrelated conditions were 0.03, 0.02, and 0.005, consistent with an improvement of the prediction score of the related compared to unrelated conditions. This finding suggests an encoding of the related context in the brain response with attention. For the unattended condition, conclusions were more ambivalent. Subject 1 showed a significant difference in prediction score (two-sided Wilcoxon signed-rank test: *W* = 2021, *p <* 0.001, Bonferroni-corrected *α* = 0.05/6) associated with a positive difference between medians: 0.01. This result is consistent with an improvement of the related condition compared to the unrelated condition. We also observed a significant difference in prediction score for subject 2 (two-sided Wilcoxon signed-rank test: *W* = 7750, *p <* 0.001, Bonferroni-corrected *α* = 0.05/6), along with a negative median difference (−0.01), consistent with an improvement of the unrelated condition over the related condition. We did not observe any significant difference between the related and unrelated context conditions for subject 3 (two-sided Wilcoxon signed-rank test, *W* = 2463, *p* = 0.0015, not significant after Bonferroni correction, *α* = 0.05/6).

In the unattended condition, the results were inconclusive, showing no clear benefit of providing related context to the LLM for generating word representations that improve brain response prediction. One possible explanation is that using too much context may introduce information that is not actually encoded in the brain when the conversation is unattended. This excessive contextual information could confuse the linear encoder, ultimately lowering prediction scores. To investigate this hypothesis, we systematically vary the context length in the next section.

### 2.3 Result 3: Contextual encoding persists without attention but at shorter context lengths

Here, we characterize the longest context length to which the model benefits in both the attended and unattended conditions. From now on, we select layer 10 for visualization because it lies well beyond the non-contextual layers while preceding the saturation or decline observed in later layers, thereby providing a representative snapshot of contextual encoding effects that are consistently observed across multiple higher layers. Results of neighboring layers from this plateau are reported in Supplementary Figure 6, as their analyses led to the same conclusions. We thus re-ran the analysis with layer 10 for different context lengths (in tokens): 1, 5, 10, 15, 50, and the full context. In Figure 5, we depict for subjects 1, 2, and 3 how the average brain score behaves as a function of context length. To ensure that the observed correlations are related to language processing, rather than being inherent to Mistral-7B’s architecture alone, we also fit an encoding model with word representations generated from Mistral-7B initialized with random weights.

**Figure 5:**
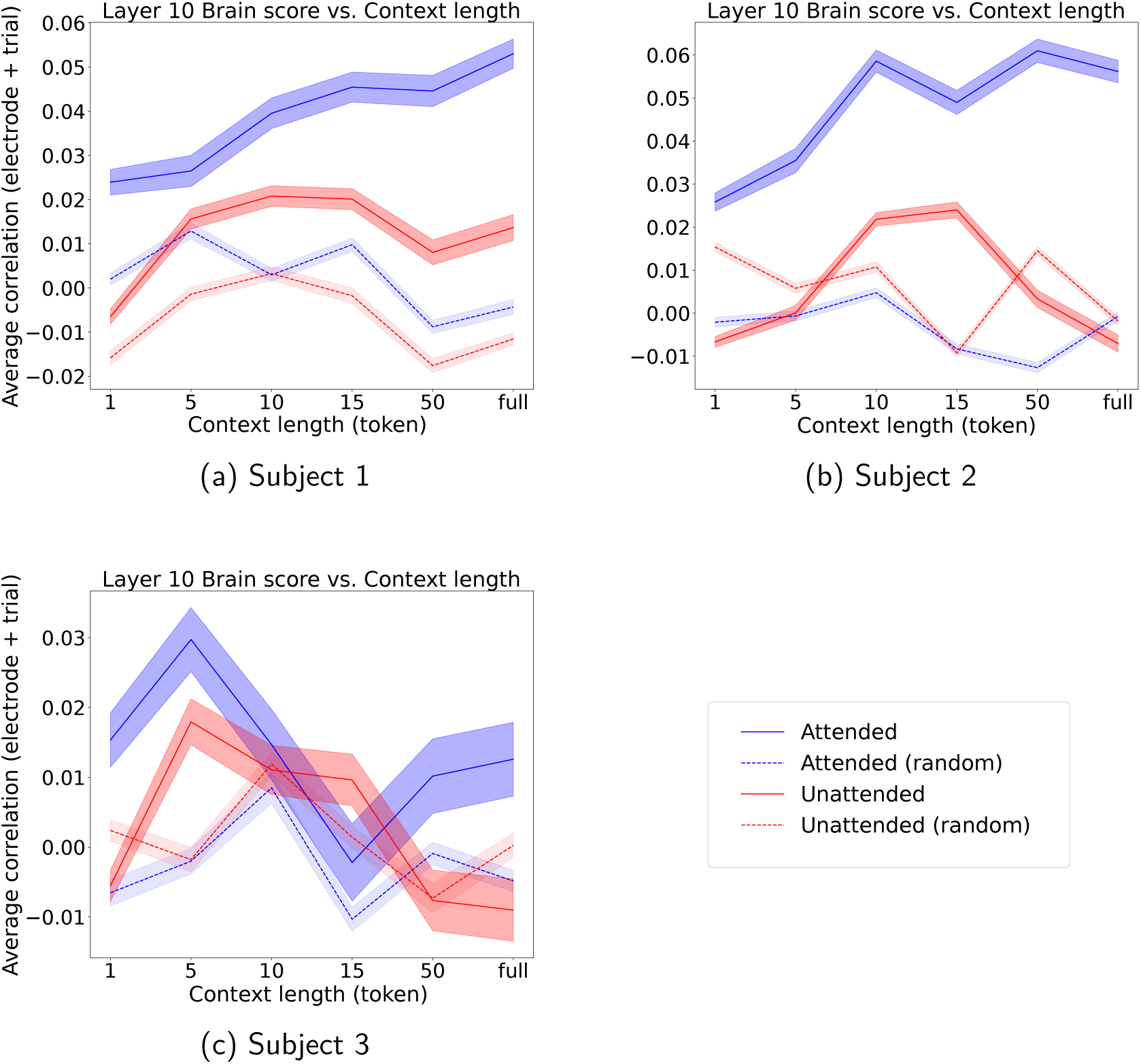
Brain prediction scores of Mistral layer 10 embeddings across context lengths. (a), (b), and (c) depict (for subjects 1, 2, and 3, respectively) the attended (blue) and unattended (red) average prediction scores across trials and electrodes using different context lengths. The blue and red dashed curves correspond to the brain prediction scores obtained with an untrained LLM. The corresponding shaded color also depicts the SEM across trial-averaged electrode prediction scores.

We observe that the average prediction score increases steadily with the context length in the attended condition, i.e., a significant difference between 15 tokens and the full context condition (two-sided Wilcoxon signed rank test, *W* = 4166, *W* = 14259, *p* < 0.001, Bonferroni-corrected *α* = 0.05/6, for subject 1 and 2, respectively) with positive median differences: 0.01 and 0.01, for subjects 1 and 2, respectively. While a statistical difference was observed between 5 and 10 tokens in subject 3 (two-sided Wilcoxon signed rank test, *W* = 1352, *p* < 0.001, Bonferroni-corrected *α* = 0.05/6), it was a decline as shown by the negative median difference (−0.02) between the two conditions.

In the unattended condition, the prediction score increase stops at 10 tokens, demonstrated by no significant difference between 10 to 15 tokens conditions (two-sided Wilcoxon signed rank test, *W* = 6186, *W* = 20889, *p* = 0.23, *p* = 0.06 for subjects 1 and 2, respectively). Then it decreased significantly from 15 to 50 tokens (two-sided Wilcoxon signed rank test, *W* = 2936, 7593, *p* < 0.001, Bonferroni-corrected *α* = 0.05/6, median differences: 0.01 and 0.03 for subjects 1 and 2, respectively). For subject 3 while a significant difference was found between 5 and 10 tokens (two-sided Wilcoxon signed rank test, *W* = 2067, *p* < 0.001, Bonferroni-corrected *α* = 0.05/6), the median difference was negative (−0.005), consistent with a decrease in prediction score between the two conditions. This finding suggests for subjects 1 and 2 that, while some context information about the unattended conversation is encoded in the brain response, its length is shorter than that of the attended conversation.

Across the three subjects, we tested the prediction score’s difference between the trained LLM and the untrained LLM condition using two-sided Wilcoxon signed-rank tests across electrodes, averaging across trials at a given context length. This analysis examines whether the encoded speech carries meaningful information beyond what is captured by an untrained model.

In Subject 1, significant differences were observed for almost all context lengths in both attended and unattended conditions (two-sided Wilcoxon signed rank test, attended: *W* = 2689 - 227, *p <* 0.001; unattended: *W* = 4705 - 2242, *p <* 0.001, Bonferroni-corrected *α* = 0.05/6), except for the attended condition at context length 5 (two-sided Wilcoxon signed rank test, *W* = 6005, *p* = 0.136). All statistical differences observed were associated with a positive median difference between the trained and untrained conditions, consistent with a higher prediction score when the LLM was trained. In Subject 2, significant differences were observed for all context lengths and both conditions (two-sided Wilcoxon signed rank test, attended: *W* = 5700 - 764, *p <* 0.001; unattended: *W* = 6937 - 15275, *p <* 0.001). While the attended case, all median differences were positive, it was only the case for the 10 and 15 tokens conditions in the unattended case (0.01 and 0.03 for 10 and 15 tokens, respectively), consistent with a higher performance for the trained LLM compared to the untrained LLM. In Subject 3, significant differences were observed for short context lengths (1 and 5) and the full context (two-sided Wilcoxon signed rank test, attended: *W* = 1366, 1049, 2277, *p <* 0.003; unattended: *W* = 1943, 1228, 1829, *p <* 0.0001). On the other hand, intermediate and long context lengths (10, 15, 50) were not significant in either attended or unattended conditions (two-sided Wilcoxon signed rank test, attended: *W* = 2988, 3043, 3214, *p* = 0.33, *p* = 0.41, *p* = 0.73; unattended: *W* = 2759, 2822, 2749, *p* = 0.10, *p* = 0.15, *p* = 0.10).

Overall, these results indicate that the trained LLM consistently outperforms the untrained LLM, supporting the conclusion that the brain encodes meaningful contextual information from attended and unattended speech embeddings.

### 2.4 Result 4: Quantifying the acoustic confounds of the LLM prediction scores

Although LLM word embeddings are derived from text rather than raw audio, they may correlate with acoustic features indirectly via the linguistic structure (e.g., lexical choice, syntactic organization). To determine whether these correlations should be interpreted as evidence of shared higher-level structure rather than acoustic encoding per se, we combine the acoustic envelope of each word with its corresponding LLM embedding in a single regression task and compare the resulting prediction score with that obtained from the acoustic envelope alone. Our hypothesis is that any substantial improvement of the LLM embedding over the speech envelope can be attributed to semantic encoding unrelated to acoustic encoding. We only show results for the first 15 layers of Mistral-7B, as subsequent layers did not improve prediction scores in previous experiments (for more details, please refer to the Discussion Section). Methodological details of the integration of the acoustic envelope into the regression task are reported in the Materials and Methods Section.

We depict the results for subjects 1, 2 and 3 in Figure 6 below.

**Figure 6:**
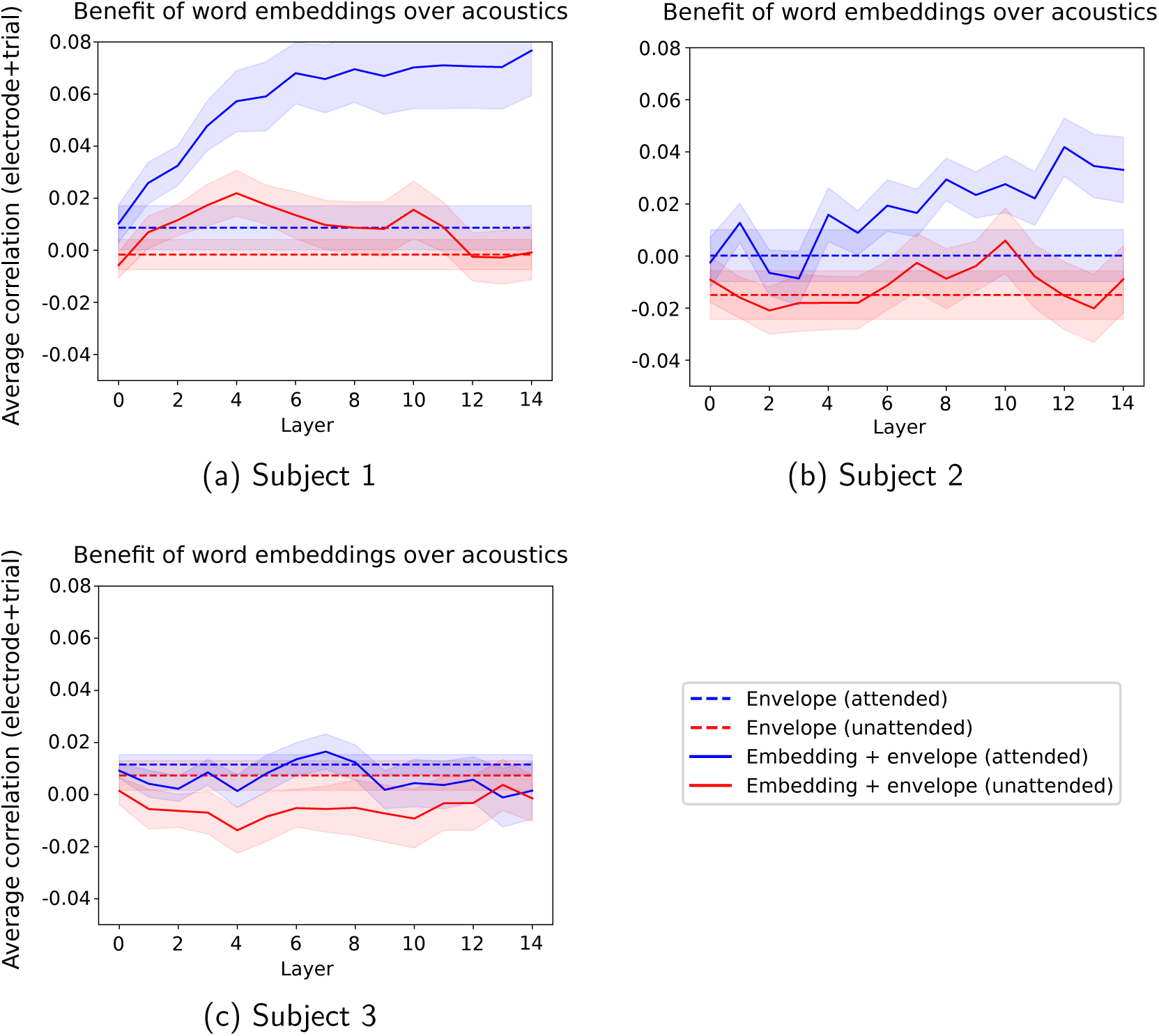
Prediction scores obtained with the speech envelope only vs. the speech envelope combined with LLM embeddings generated with the full context for each word. (a), (b), and (c) depict (for subjects 1, 2, and 3, respectively) the attended (continuous blue) and unattended (continuous red) average prediction scores across trials and electrodes for the combination of LLM embeddings generated with the maximal context for each word and the corresponding acoustic envelope. The blue and red dashed lines correspond to the prediction scores obtained only with the attended and unattended acoustic envelopes, respectively. The corresponding shaded color also depicts the SEM across trial-averaged electrode prediction scores.

For subjects 1 and 2, we see a significant benefit to the prediction score of the LLM embeddings beyond the acoustic envelope for the attended stream (two-sided Wilcoxon signed-rank test, subject 1: *W* = 0, *p <* 0.001; subject 2: *W* = 7.0, *p* = 0.0011, Bonferroni-corrected *α* = 0.05/6). For subject 3, we found the opposite effect (two-sided Wilcoxon signed-rank test, *W* = 9, *p <* 0.001, Bonferroni-corrected *α* = 0.05/6). This aligns with previous results showing no improvement in prediction scores across layers, possibly due to poor electrode coverage and responsiveness. We thus ignore subject 3 for the rest of this subsection.

For the unattended stream, the conclusions were more mixed when considering the maximal context for each word. For subject 1, a benefit of the LLM embeddings beyond acoustics was observed (two-sided Wilcoxon signed-rank test, *W* = 9, *p <* 0.01, Bonferroni-corrected *α* = 0.05/6), while not for subject 2 (two-sided Wilcoxon signed-rank test, *W* = 31, *p* = 0.11).

Overall, we note that prediction scores obtained using the acoustic envelope alone are generally low, reflecting the relatively small number of electrodes with high scores compared with the model integrating LLM embeddings (see Supplementary Figure 8). This is in line with the minimal speech-responsive electrode selection that was made previously with this dataset (Choudhari et al., 2024), and gives compelling evidence that LLM embeddings not only carry semantic information beyond acoustics on average, but also lead to higher prediction scores for more electrodes. These results confirm that the prediction scores estimated in the previous experiments for subjects 1 and 2 stem from higher-level speech features than mere acoustic encoding when considering the attended stream. As hypothesized previously, the context length encoded for the unattended stream may be shorter (see Figure 5).

We thus depict, for the unattended stream, the benefit of LLM embeddings beyond acoustics, using a shorter context length of 10 tokens for subjects 1 and 2. We chose 10 tokens because it is the length at which the prediction score saturates in Figure 5 for subjects 1 and 2. We depict the results in Figure 7 below:

**Figure 7:**
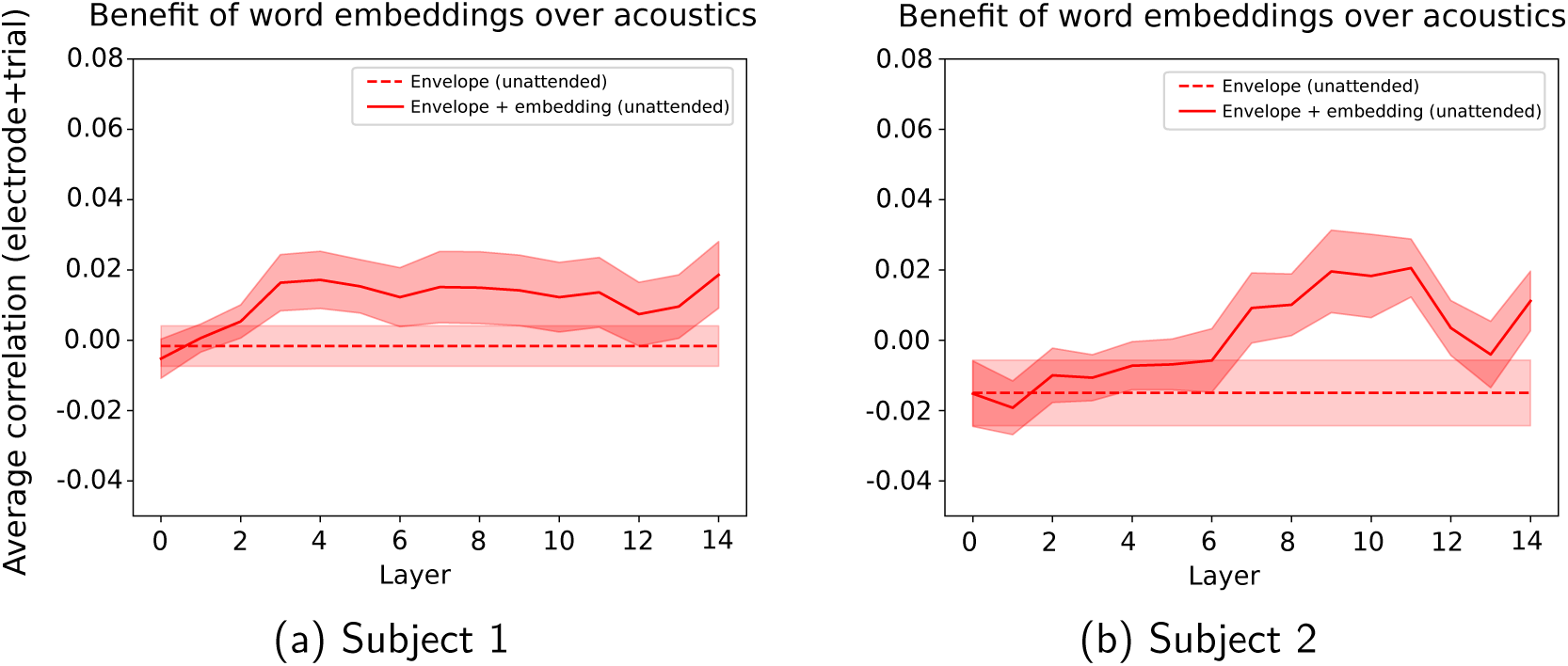
Unattended stream prediction scores obtained with the speech envelope only vs. the speech envelope combined with LLM embeddings generated with the a shorter context length (10 tokens) for each word. (a) and (b) depict (for subjects 1 and 2, respectively) the unattended (continuous red) average prediction scores across trials and electrodes for the combination of LLM embeddings generated with a 10-tokens context length for each word and the corresponding acoustic envelope. The red dashed line corresponds to the prediction scores obtained with the unattended acoustic envelope only. The corresponding shaded color also depicts the SEM across trial-averaged electrode prediction scores.

We observe a significant increase in the prediction score when adding LLM embeddings to the acoustic envelope at a context length of 10 tokens for subjects 1 and 2 (two-sided Wilcoxon signed-rank test, subject 1: *W* = 2.0, *p <* 0.001; subject 2: *W* = 3.0, *p <* 0.001, Bonferroni-corrected *α* = 0.05/6). These results confirm that, at shorter context lengths, the prediction scores estimated in the previous experiments for subjects 1 and 2 are driven by higher-level speech features rather than by mere acoustic encoding when considering the unattended stream.

## 3 Discussion

In this study, we explored the extent to which contextual LLM-based linguistic features can predict brain responses to speech in a multi-talker listening environment beyond acoustics. Furthermore, we characterized the range of context encoded in the brain when a conversation is attended or unattended.

In Result 1, we showed that the LLM gradually learns contextual information when increasing the layer index, and we found that this improved the brain score until a plateau is reached. The performance then decreases, potentially because Mistral-7B is a generative LLM, which means it will output an answer to a text input. In late layers, the embeddings may thus incorporate more information related to the generated response and unrelated to the input prompt, which could explain the decrease in the brain score in later layers. We found this for the attended condition but also, in the case of subject 1, for the unattended condition. Although prediction scores reach a lower-magnitude plateau, this suggests contextual tracking without attention. Anderson et al., 2024 performed a similar analysis combining features extracted from an ASR model (i.e., Whisper), the word surprisal from an LLM (i.e., GPT-2), and the speech envelope with its first derivative. They demonstrated that later Whisper layers were more accurate EEG predictors for the attended stream, providing an added value over the speech envelope and its first derivative. Moreover, they found a unique contribution of the word surprisal, and the speech envelope with its derivative to predict the response measured by certain electrodes. This suggests a non-contextual and contextual tracking of the attended stream. On the other hand, the unique contribution of the word surprisal and of Whisper deep layers disappeared for the unattended stream, suggesting the absence of contextual information encoded in the brain.

Additionally, an earlier EEG study using linear models demonstrated no lexical processing when a speaker was unattended (Brodbeck et al., 2018). We explain our different conclusions with the low SNR of EEG recordings. Multiple studies demonstrated the ability of LLM-generated features to be good predictors of the brain response in single-talker paradigms, notably with fMRI (Antonello et al., 2023; Caucheteux et al., 2023) and ECoG (Goldstein et al., 2022; Mischler et al., 2024). Our current finding expands the understanding of contextual encoding in a multi-talker scenario, by suggesting that some contextual linguistic information from the unattended speaker is being processed in the brain, and on a shorter time scale than for the attended speaker. This is the first time it is done with LLM-generated features, and it also questions earlier EEG findings, setting the limits of such recording modality to estimate the processing of linguistic features.

In Result 2, we showed that the model relies on the correct contextual information to improve predictions of brain responses. To test this, we compared prediction scores across layers when Mistral-7B was fed the original trial context for each word versus an unrelated context taken from another story (“the black willow”). In the attended case, the unrelated context condition led to lower prediction scores across layers, underscoring the encoding of context information beyond the embedding of the word of interest. In the unattended case, word representations may have included information from prior words that were not actually encoded in the brain. This hypothesis was further explored and supported in Result 3.

Taken together, the layer-wise analysis (Result 1) and the context-manipulation experiment (Result 2) suggest that the predictive power of LLM embeddings does not arise from lexical or part-of-speech information alone, but from representations that depend on preceding linguistic context. Because the identity and syntactic category of the target word were preserved while the surrounding context was altered, the observed drop in prediction scores indicates sensitivity to context-dependent representations. This contextual sensitivity plausibly reflects semantic integration processes, extending beyond shallow syntactic features or low-level acoustic tracking.

In Result 3, we explored multiple context lengths to characterize whether the attended and unattended context encoding were in the same length range. We found that for subjects 1 and 2, the unattended context length tracked reached a plateau earlier than that of the attended. Varying the context length provided to models has been done in multiple ECoG studies for single-speaker experiments with LLMs (Goldstein et al., 2022; Mischler et al., 2024), also demonstrating a gradual increase of the prediction scores when increasing the context length. A recent EEG study (Anderson et al., 2024) investigated how ASR-extracted features relate to the brain response in a multi-talker environment. In line with previous EEG findings (Brodbeck et al., 2018), the authors did not find any linguistic contribution for the unattended stream, which contradicts our current findings for subjects 1 and 2. We explain this divergence with the low SNR of EEG, leading to minimal contributions of linguistic features to the EEG response, even in single-speaker paradigms compared to acoustics or lexical segmentation features (Gillis et al., 2021; Puffay et al., 2023). For subject 3, we did not find the same context length tracking for both the attended and unattended conditions. Two potential explanations are: (1) the number of electrodes is minimal in the frontal lobe (see Supplementary Figure 5a), which contains areas encoding context (Caucheteux et al., 2023), and (2) the electrodes are less responsive, leading to poorer prediction scores than in subjects 1 and 2 (see Supplementary Figures 5c-e).

In Result 4, we quantified the added value of LLM-generated word embeddings over their acoustic envelope, to ensure that the prediction scores obtained with embeddings were not only reflecting acoustic processing, but also semantic. We found a significant benefit of LLM embeddings beyond acoustics for both the attended and unattended streams, with a shorter context length encoding for the unattended stream, which confirmed our findings that non-acoustic linguistic processing happened with and without attention. Interestingly, the prediction score values were almost identical between the embeddings from layer 0 and the acoustic envelope. Considering that the speech envelope and Mistral-7B’s layer 0 carry information about the word of interest only, this reinforces the claim that the prediction score increase observed across layers may reflect contextual processing. This result aligns with a previous study (Caucheteux et al., 2021), using embeddings from layer 0 as non-contextual lexical controls.

First and foremost, although ECoG and sEEG provide highly localized brain responses with a high signal-to-noise ratio (SNR), our study includes only three subjects, making it difficult to draw general conclusions. Additionally, subject 3 did not exhibit the same pattern, potentially due to limited electrode coverage or reduced responsiveness. Nonetheless, the results reveal a compelling pattern regarding contextual tracking across attentional states that warrants further investigation.

The limited electrode coverage also limited our ability to identify neuro-anatomical patterns related to the impact of attention on context. To have a homogeneous coverage across subjects, and if fast brain responses are not of interest, fMRI might offer an alternative to investigate neuro-anatomical insights using LLMs (e.g., Antonello et al., 2023; Caucheteux et al., 2023).

Besides discrepancies in electrode coverage between subjects, attention might not be a very localized phenomenon. We can think of attention as operating on different levels, such as acoustic attention, (sub-)lexical attention, and semantic attention. As a result, various brain areas involved in speech processing may be influenced by attention.

However, our text-based features lack acoustic details like tone, accent, and volume. While focusing solely on (sub-)lexical and linguistic information offers valuable insights into how these aspects are encoded, some brain regions might primarily respond to acoustic features of speech (O’Sullivan et al., 2019), limiting our ability to capture localized attention effects. A potential solution could be to combine this analysis with Automatic Speech Recognition (ASR) models like Whisper (Anderson et al., 2024; Radford et al., 2022), allowing us to incorporate acoustic information while adjusting for context in the word representations.

Lastly, the behavioral data analysis from Choudhari et al., 2024 indicated that the uncued conversation might have been attended occasionally (up to 8% of repeated word detection for subject 1). To alleviate the impact of these attention drops on our conclusions, we excluded the trials containing such events and showed that contextual tracking of the unattended stream remains (See Supplementary Figure 4). Our word detection task occurs every 7 s, thus not involving every word from the trials. Consequently, not every attention drop might be detected in our behavioral measurement. An objective measurement of auditory attention for every segment, such as alpha activity (Deng et al., 2020) or the performance of an auditory attention decoding model, could further reinforce our claim. On average across our three subjects, Choudhari et al., 2024 demonstrated a high auditory attention decoding accuracy (e.g., 82% using 2 s segments), suggesting minimal attention drops.

In conclusion, this study presents significant insights into how contextual linguistic features from LLMs, specifically Mistral-7B, can predict ECoG responses to speech in a multi-talker environment. By characterizing the contextual information encoded across attentional states, we gained a deeper understanding of auditory attention, which had previously ignored the contextual component of linguistic tracking. Our modeling approach could be applied to perform auditory attention decoding, which usually uses acoustic speech features, or to further explore the neural mechanisms of attention with a broader electrode coverage and more participants.

On a broader scope, characterizing context encoding could be used to measure speech understanding objectively. One could, for instance, imagine evaluating the context encoding when listening to a foreign language (Bhaya-Grossman et al., 2025) or degraded speech.

## 4 Methods and Materials

### 4.1 Subjects and neural data

We used data from a previous study (Choudhari et al., 2024), including 3 subjects, of whom one (subject 1) was from Irving Medical Center at Columbia University, and two (subjects 2 and 3) were from North Shore University Hospital. All participants were undergoing clinical treatment for epilepsy and provided informed consent per the local Institutional Review Board (IRB) regulations. Neural data were collected as subjects performed the task with ECoG. Subject 1 only had sEEG depth electrodes implanted over their left brain hemisphere. Subjects 2 and 3 had sEEG depth electrodes as well as subdural ECoG grid electrodes implanted over the left hemispheres of their brains (see Figure 2).

Subject 1’s neural data were recorded using Natus Quantum hardware at a sampling rate of 1024 kHz, and subjects 2 and 3’s neural data were recorded using Tucker-Davis Technologies (TDT) hardware at a sampling rate of 1526 Hz. The left and right audio channels played to participants were also recorded in sync with neural signals to facilitate segmenting neural data into trials for offline analysis. The trials were spatialized using head-related transfer functions (HRTFs) and delivered to the subjects via earphones.

### 4.2 Neural data preprocessing

Following Choudhari et al., 2024, neural data were first resampled to 1000 Hz, then a common average reference was performed to reduce recording noise. The neural data were then further downsampled to 400 Hz. Line noise at 60 Hz and its harmonics (up to 180 Hz) were removed using a notch filter. A 1000th-order finite impulse response (FIR) notch filter was designed using MATLAB’s *fir2* function, and zero-phase filtering was achieved with *filtfilt*. To extract the envelope of the high gamma band (70 – 150 Hz), the neural data were first filtered with eight Gaussian filters, each with a width of 10 Hz, spaced consecutively between 70 and 150 Hz. The envelopes of the outputs of these filters were obtained by computing the absolute value of their Hilbert transform. The final envelope of the high gamma band was obtained by computing the mean of the individual envelopes yielded by the eight filters and further downsampling to 100 Hz.

### 4.3 Experiment Design

The experiment included 25 multi-talker trials, each with an average duration of 44.2 seconds (SD = 2.0 seconds), and 138 words on average (SD=6.86). As shown in Figure 1 (a), each trial featured two simultaneous and independent conversations, which were spatially separated and moved continuously within the frontal half of the subject’s horizontal plane (azimuth range: −90 ° to +90 °), occasionally crossing each other. The intensity of both conversations remained equal and constant throughout the experiment; the participants adjusted the overall mixture volume to a comfortable level at the beginning of the task. Both conversations had the same power (RMS); all talkers were native American English speakers. Diotic background noise (Barker et al., 2015; Spahr et al., 2012) — either non-stationary noise (i.e., street noise, people chatter, honking) or stationary noise (babble)— was mixed in with the conversations at a power 9 or 12 dB lower than the conversation stream (25% of trials with babble −9 dB, 25% with babble −12 dB, 25% with pedestrian −9 dB and 25% with pedestrian −12 dB).

In these conversations, a total of 8 different talkers (4 male, 4 female) alternated speaking. The assignment of genders to conversations were balanced to avoid bias (i.e., 25% male-male, 25% female-female, 25% female-male, and 25% male-female). As illustrated in Figure 1 (b), in the attended conversation, a talker switch occurred at the 50% mark of the trial duration. For the unattended conversation, two talker switches happened—one around the 25% mark and the other near the 75% mark. Participants were instructed at the start of each trial to attend to the conversation that began first and to maintain attention on that conversation throughout the trial, regardless of talker switches. They were told to press a button whenever they detected a repeated word in the attended conversation. Talker switches were not explicitly cued to participants; they were expected to follow the continuity of the conversation rather than a specific talker’s voice (Choudhari et al., 2024).

To ensure that subjects remained focused on the attended conversation and control for it behaviorally, a 1-back detection task was included in the experiment. Both conversations included occasional repetition of words, strategically chosen to avoid temporal overlap between the two conversation streams. A repeated word was created by duplicating the talker’s voice waveform for that word. On average, the interval between the onset of two repeated words within a conversation was 7.0 seconds (SD = 1.0 seconds). The assignment of male and female talkers to different segments of the conversations was counterbalanced across trials to ensure equal durations of concurrent conversations with either the same or different genders.

### 4.4 Behavioral data

The push-button responses of subjects to repeated words in the conversation they were following were used to determine which conversation the subject was attending. A repeated word was considered correctly detected only if a button press occurred within two seconds of its onset. As reported in Choudhari et al., 2024, 67%, 75%, and 73% of the repeated words in the cued (attended) conversation were correctly detected for subjects 1, 2, and 3, respectively. In contrast, only 8%, 6%, and 1% of the repeated words in the uncued (unattended) conversation were correctly detected, suggesting that there were instances when subjects may have been attending to the unattended conversation as well.

### 4.5 Large language models to generate word representations

We conducted our analyses using Mistral-7B, a decoder-only transformer model comprising an initial token-embedding layer followed by 32 transformer blocks, as described in the architecture and pre-training overview (Jiang et al., 2023). Mistral-7B yielded the highest brain response prediction scores in a study comparing different open-source LLMs for generating word representations used as ECoG response predictors (Mischler et al., 2024). Importantly, the initial token-embedding layer does not encode positional or contextual information; we refer to it as layer 0 in this study. We downloaded the model from the Hugging Face platform.

Embeddings extracted from Mistral-7B are interpreted as model-internal representations of word meaning that progressively integrate linguistic context across layers. While the initial token embedding layer (layer 0) reflects purely lexical information, successive transformer layers incorporate increasing amounts of contextual information from preceding tokens via self-attention. We deliberately used a pretrained, task-agnostic language model without task-specific fine-tuning, such that the extracted embeddings reflect general-purpose statistical regularities of natural language rather than representations optimized for the experimental paradigm. This design allows us to investigate how neural activity tracks contextual and non-contextual language representations during naturalistic speech perception, without introducing task-dependent biases.

For each trial of both the attended and unattended speech streams, we fed the corresponding transcripts to Mistral-7B and extracted the embeddings of each layer. For each trial, the resulting dimensions were, therefore, *T* × *D* × *L*, with *T* the number of tokens, *D* the dimension of each embedding (i.e., 4096 for Mistral-7B), and *L* the number of layers (i.e., 32). For multi-token words, we selected the final token embedding to represent the word, as it captures the most complete contextual information. For experiments that varied the context provided to Mistral, we truncated the attention mask to include only the previous *M* tokens, *M* being the defined context length.

### 4.6 Word envelopes as controls for acoustic encoding

To control for acoustic information that can be derived from LLM embeddings, we extracted the speech envelope for each word. To do so, we first generated the auditory spectrogram of the raw audio using the naplib-python package. The model used to generate the auditory spectrogram includes a cochlear filter bank of 128 logarithmically-spaced constant-Q filters, a hair cell model with a low-pass filter and a nonlinear compression function, and a lateral inhibitory network along the spectral axis. The envelope of each frequency band gives the time-frequency representation. To obtain the acoustic envelope, we averaged the auditory spectrogram over the frequency axis to obtain a one-dimensional speech envelope. For each word, we then extracted a lagged-envelope matrix consisting of 7 windows. The first window starts at the word onset and ends at the word length median (i.e., 250 ms); six lags are then extracted, with a stride of 50 ms. We selected the median word length as a reasonable fixed window length for most words, since our regression models do not accept variable-length inputs.

### 4.7 Ridge regression mapping from word embeddings to neural responses

To evaluate how predictive of different brain responses the LLM word representations are, we implemented ridge regressions to map the different layer embeddings of Mistral to different electrode responses in the brain of our subjects (see Figure 2). We selected 200 ms windows for each electrode to capture word responses, considering seven different lags starting at word onset, with a stride of 50 ms, thus covering a 500 ms span post-word-onset. We temporally averaged neural responses over 200 ms frames and used ridge regression to predict (averaged) neural responses from model embeddings as a function of lag. For each trial transcript, layer-specific word representations were computed with varying context lengths specified further.

Thus, for each trial and electrode, we fitted a ridge regression model with leave-one-trial-out cross-validation to predict the average lagged word responses from the embeddings of each layer of Mistral-7B. The decision to exclude a trial reduces the risk of context leakage, which the model could learn instead of the actual response-language relationship. For each training fold, we swept over a range of regularization parameters (log space between 10^−2^ and 10^4^). Although there were small variations in brain reconstruction scores (see Supplementary Figure 7), we combined the seven lags in our regression models to allow them to associate the word embeddings with different brain response delays.

In subsection 2.4, we controlled for the encoding of acoustic information by combining the word embeddings with the corresponding acoustic envelope. For each word, we concatenated the LLM embedding and the lagged envelopes to obtain an input per word of dimension *D* + *nT*, with *D* = 4096 being the dimensions of Mistral-7B embeddings, *n* = 7 being the number of lags, and *T* = 250 being the number of time samples for each envelope window.

### 4.8 Context modulation

We manipulated the transcript multiple times before word embedding generation using Mistral-7B.

In Result 2, we compared the brain reconstruction scores using word representations with the original context presented to the participants to the word representations generated using an unrelated context. This experiment aims to show that context contributes to the brain reconstruction score, suggesting an encoding of contextual information in the brain. To generate word representations with an unrelated context, we identified for each word *W* the number of words *K* before it within the trial. We then selected *K* consecutive words from a different transcript (”the black willow”, from the MAG-MESC dataset Gwilliams et al., 2023), adding *W* at the end of the word’s sequence. We further fed this context to Mistral-7B to generate the representation of *W* with an unrelated context.

In Result 3, we varied the number of tokens fed to Mistral before each token of interest to investigate the impact of the context length used for word representation generation on the brain reconstruction scores. We tested the following lengths (in tokens): 1, 5, 10, 50, for both the attended and unattended conditions.

## Data and Code Availability

The data that support the findings of this study are available on request from the corresponding author (N.M.). The data are not publicly available due to privacy or ethical restrictions. The code base to run our analyses is available on GitHub: https://github.com/corentinpuffay/LLM_ECoG.

## Author Contributions

Vishal Choudhari (V.C.), Stephan Bickel (S.B.), Ashesh D. Mehta (A.D.M.), Catherine Schevon (C.S.) and Guy M. McKhann (G.M.M) contributed to data collection. Gavin Mischler Moore (G.M.M.), and V.C. contributed to conceptualization, methodology, and revision of the manuscript. Corentin Puffay (C.P.) contributed to the conceptualization, methodology, data analysis, and writing of the manuscript. Jonas Vanthornhout (J.V.) contributed to conceptualization, methodology, and revision of the manuscript. Tom Francart (T.F.) contributed to the revision of the manuscript. Nima Mesgarani (N.M.) supervised the project and contributed to conceptualization and methodology.

## Acknowledgements

Sukru Samet Dindar for helping me familiarize myself with the dataset. Richard Antonello for the useful discussions and for providing the code base for the ridge regression optimization.

## Supplementary Material

See supplementary_material.tex.

