## Supplementary figures for "Large Language Models Reveal the Neural Tracking of Linguistic Context in Attended and Unattended Multi-Talker Speech"

February 24, 2026

### 1 Text S1: Impact of repeated words on the prediction scores

The experimental design includes repeated words in each trial, which may affect the word embeddings generated by the LLM and, in turn, impact the correlations obtained when predicting brain responses. To address this, we excluded the second occurrence of repeated words from the transcripts, regenerated the word embeddings, and repeated the analysis. The results for subjects 1, 2, and 3 are shown in

Supplementary Figure 2. No differences that would affect our conclusions were observed.

### 2 Text S2: Impact of speaker ID on the prediction scores

We observed a significant increase in correlation for the unattended transcript in subject 1. Some brain regions are known to respond to individual speakers, and we hypothesized that this increase might come from an electrode response to a specific speaker, rather than contextual information from all unattended speakers. To ensure this finding is not due to a spurious speaker identification, we isolate and average the trials where speaker 0 is present, then repeat the process for speaker 1, and so on for all speakers. This results in eight curves, one for each speaker. The resulting correlations are shown in Supplementary Figure 3. Besides speaker 2, we observe that all speakers contribute to the observed correlation increase, ruling out the possibility that the increase is due to a specific speaker.

### 3 Text S3: Impact of attention drops on the prediction scores

To measure attention behaviorally, the participants performed a word detection task during the experiment. Attention drops were considered when a button press occurred within 2 s after the second occurrence of a repeated word in the unattended conversation. These attention drops occurred for subjects 1, 2 and 3, in 6, 2 and 1 trials respectively. To verify that these attention drops are not at the origin of the semantic tracking in the unattended stream, we excluded the above-mentioned trials from the analyses. We observed no changes in the results (see Supplementary Figure 4), and therefore assume that behavioral attention drops are not at the origin of the observed semantic tracking.

### 4 Supplementary Figures

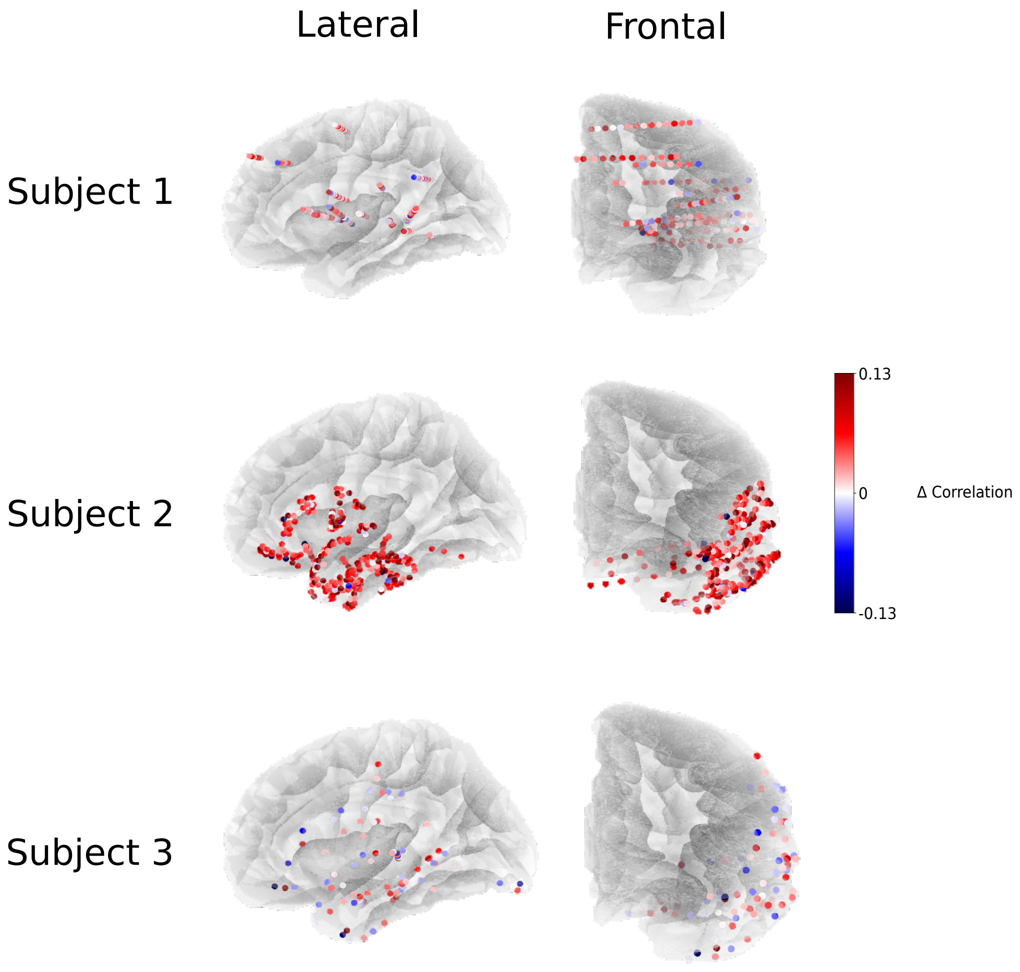


Figure 1: **Brain mapping of prediction scores' difference between attended and unattended conditions (layer 10).**

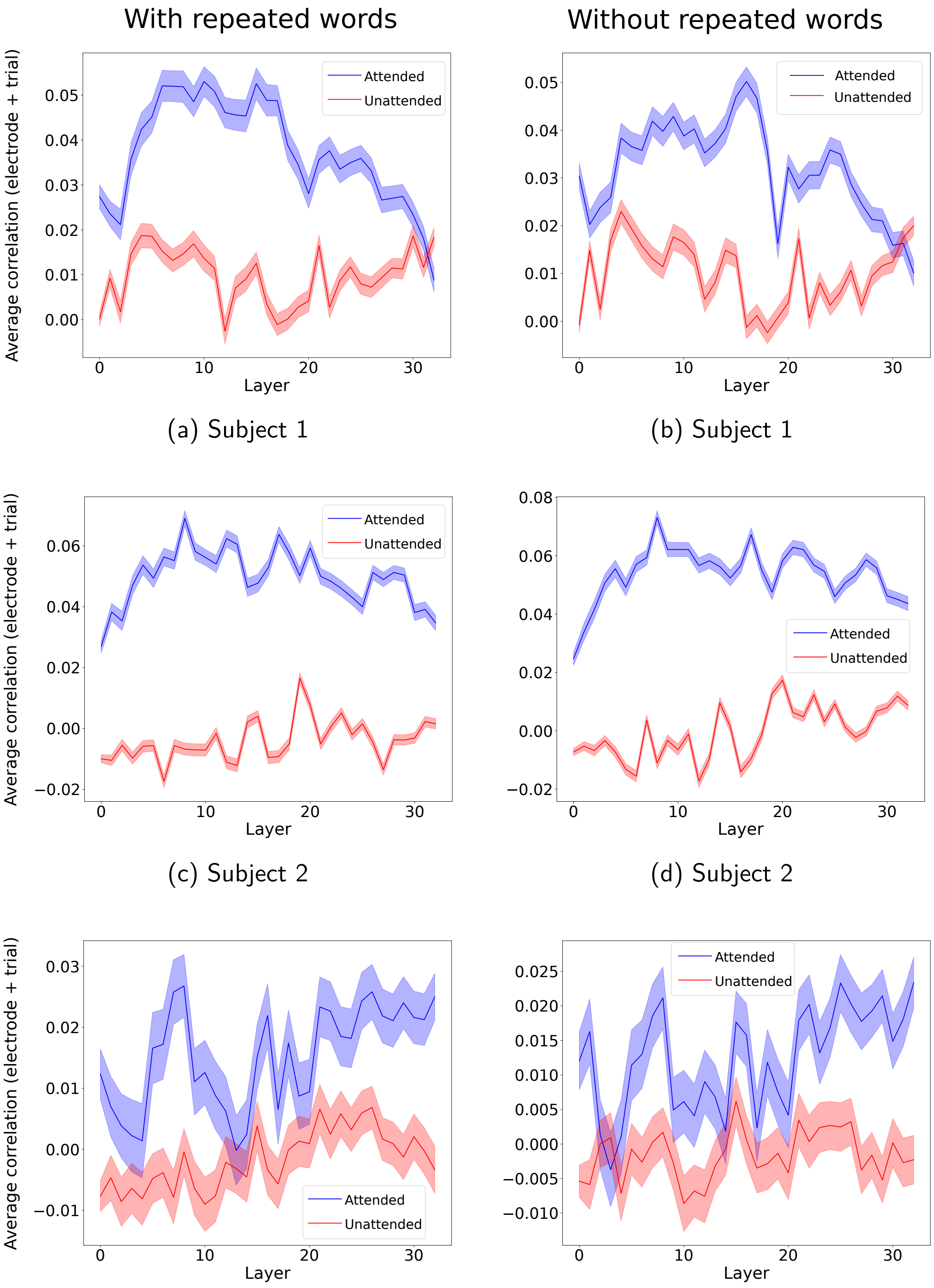


(a) Subject 1 (b) Subject 1

(c) Subject 2 (d) Subject 2

(e) Subject 3 (f) Subject 3

Figure 2: **Impact of repeated words on brain prediction scores of LLM word embeddings across layers with attention modulation.**

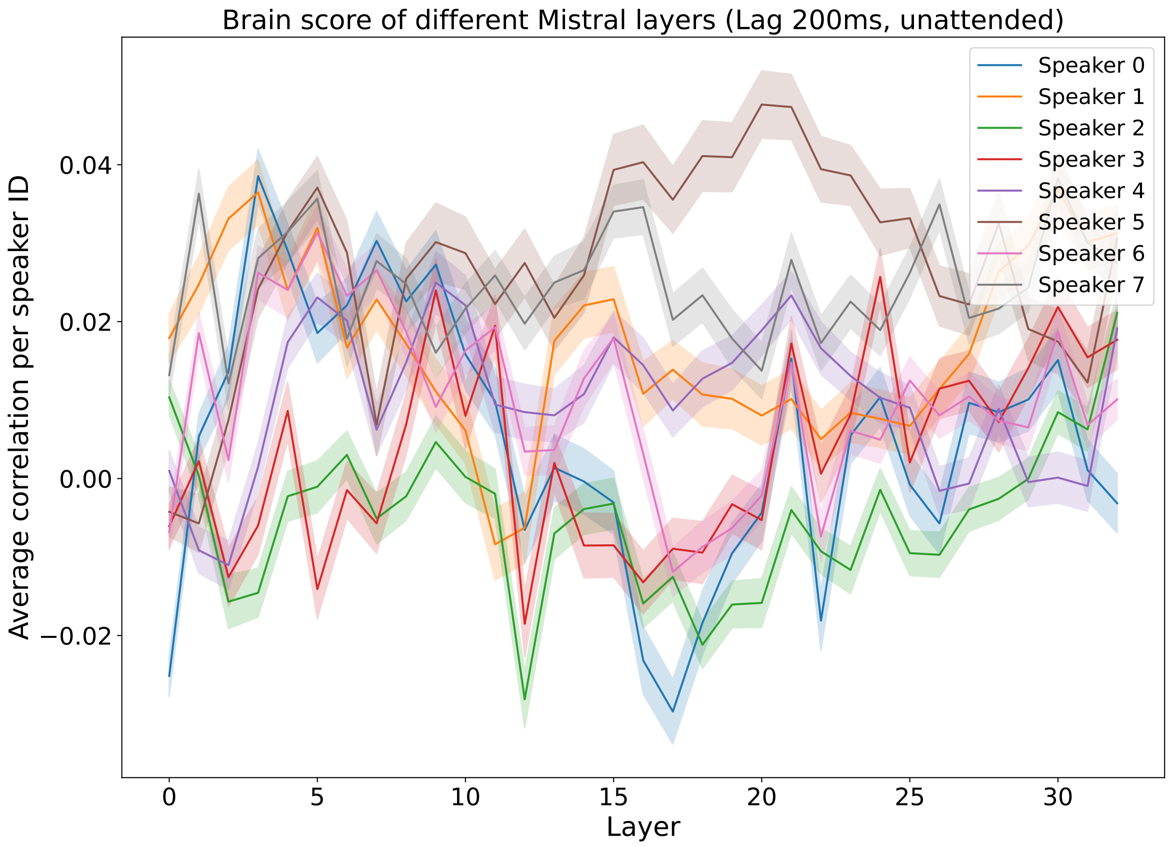


Figure 3: **Average brain prediction score for each speaker across Mistral-7B layers for the unattended transcript in subject 1.** We isolated and averaged the trials where speaker 0 is present, then repeat the process for speaker 1, and so on for speakers 0–7. This results in eight curves, one for each speaker. Selected window: $[WO + 200ms, WO + 400ms]$, $WO$ being the word onset timing.

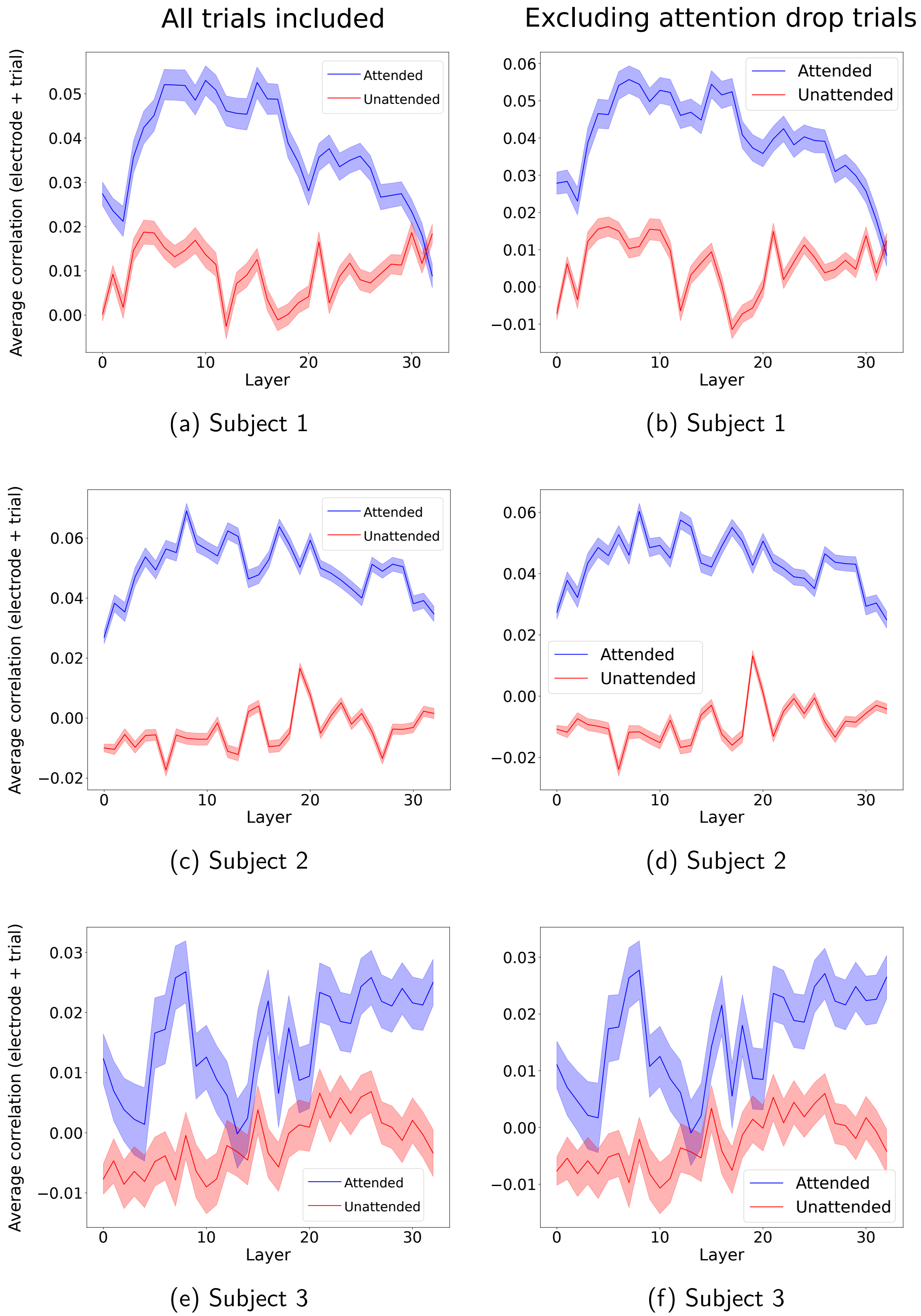


(a) Subject 1 (b) Subject 1

(c) Subject 2 (d) Subject 2

(e) Subject 3 (f) Subject 3

Figure 4: **Impact of attention drops measured behaviorally on brain prediction scores of LLM word embeddings across layers with attention modulation.**

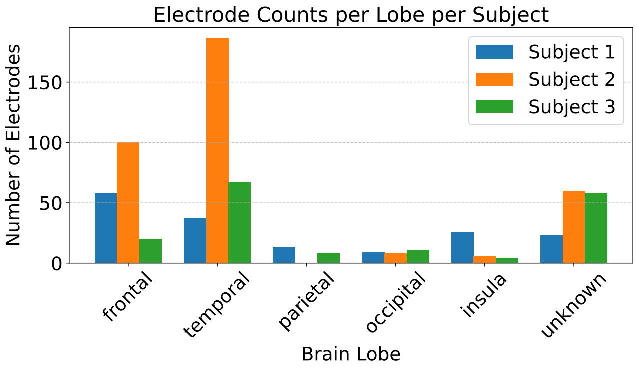


(a) Electrode counts per lobe per subject

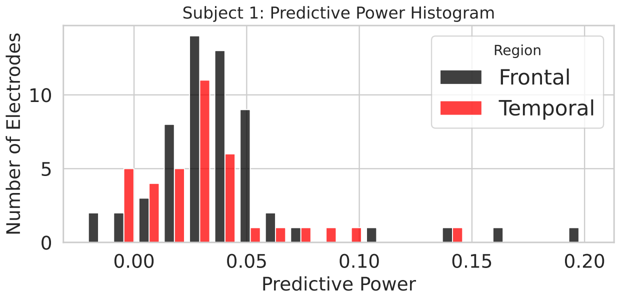


(b) Subject 1

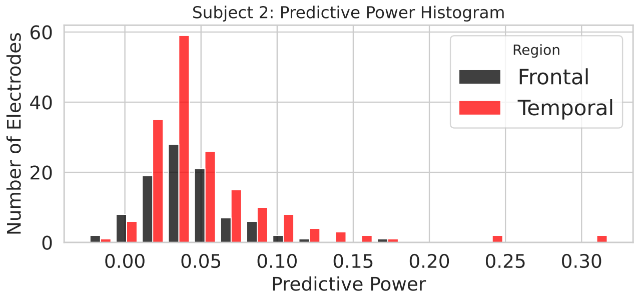


(c) Subject 2

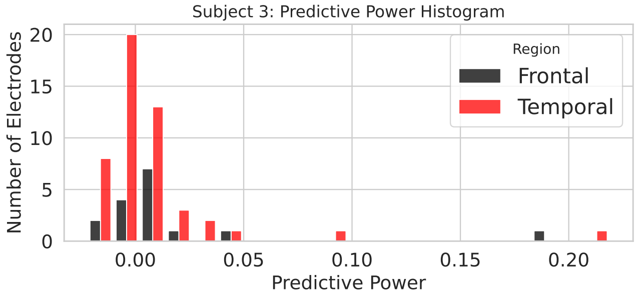


(d) Subject 3

Figure 5: **Lobe distribution of electrode per subject and histograms of trial-averaged predictive scores per electrode from the frontal and temporal lobe.**

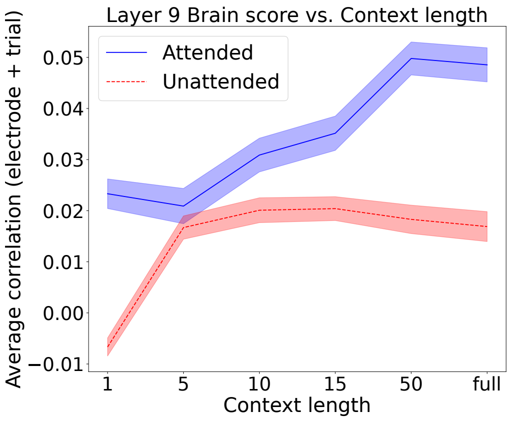


(a) Subject 1

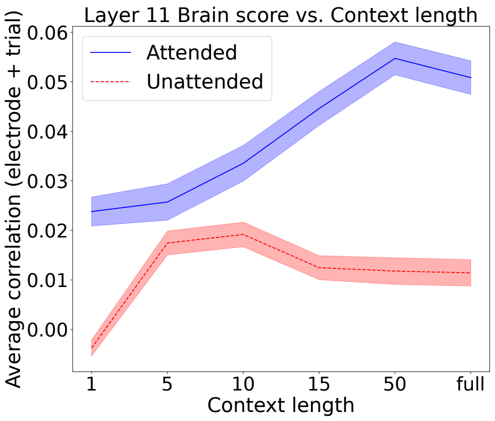


(b) Subject 1

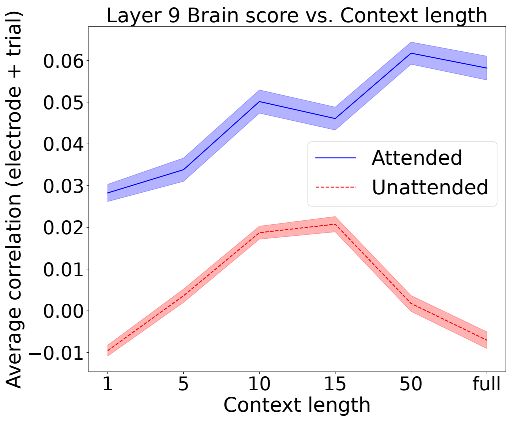


(c) Subject 2

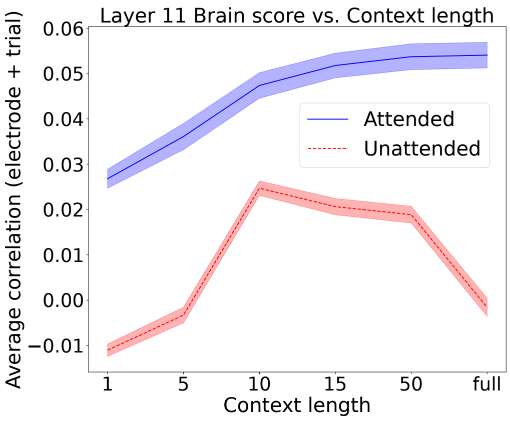


(d) Subject 2

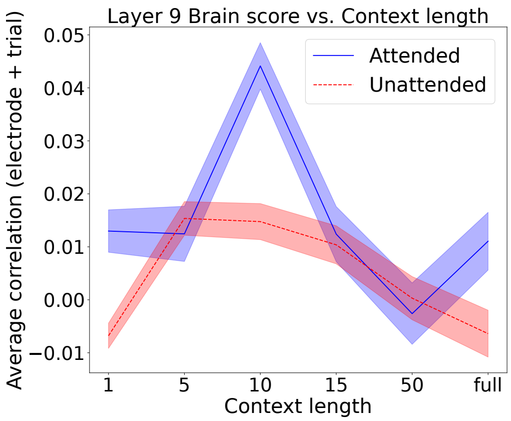


(e) Subject 3

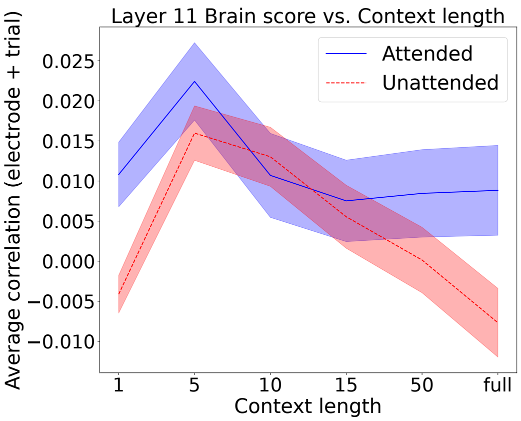


(f) Subject 3

Figure 6: **Brain prediction scores of Mistral layer 9 and 11 embeddings across context lengths.**

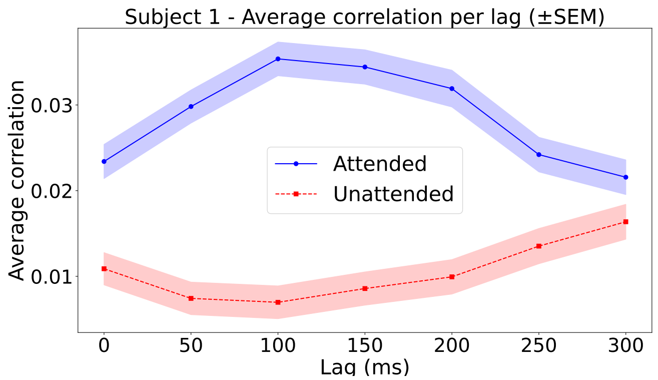


(a) Subject 1

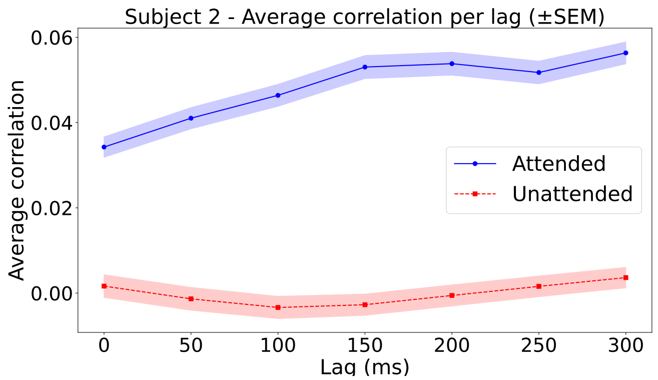


(b) Subject 2

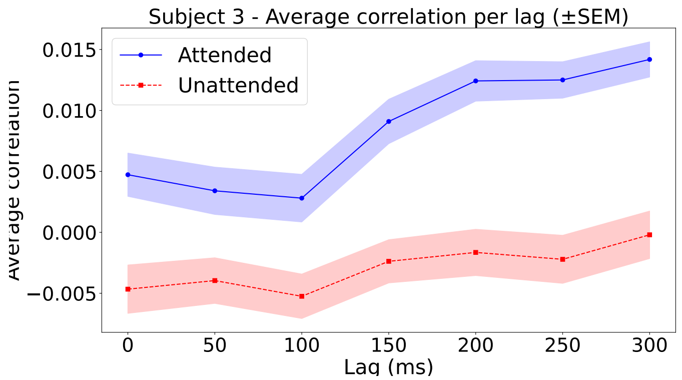


(c) Subject 3

Figure 7: **Average brain reconstruction score across post-word onset lags.** The average brain reconstruction scores are averaged across electrodes, trials and layers for different lags. Lag X means that the 200 ms window selected starts from X ms after the word onset.

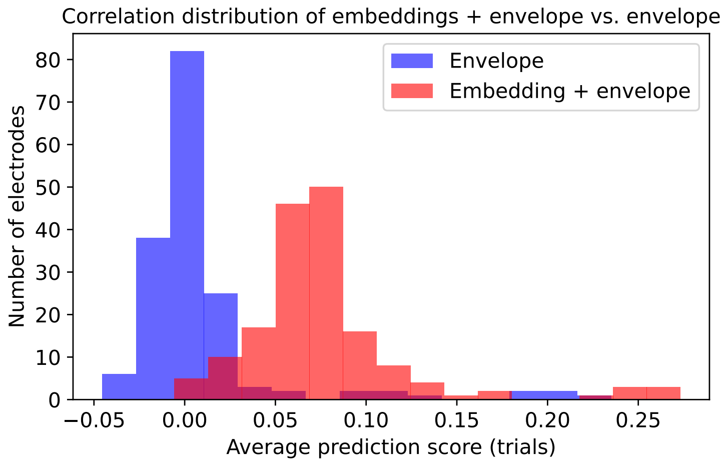


(a) Subject 1

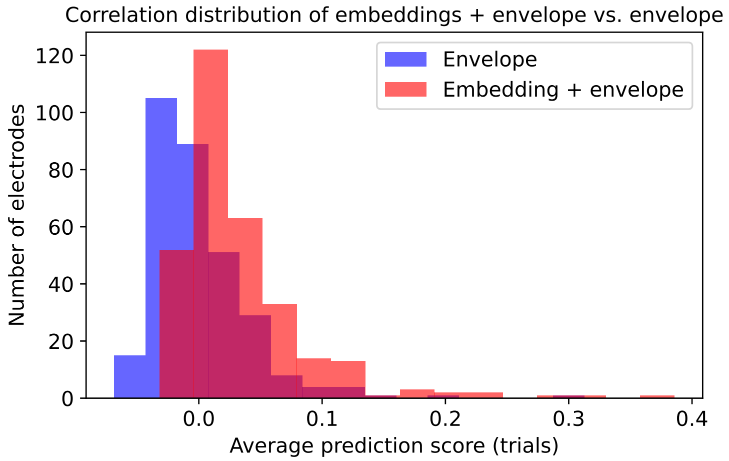


(b) Subject 2

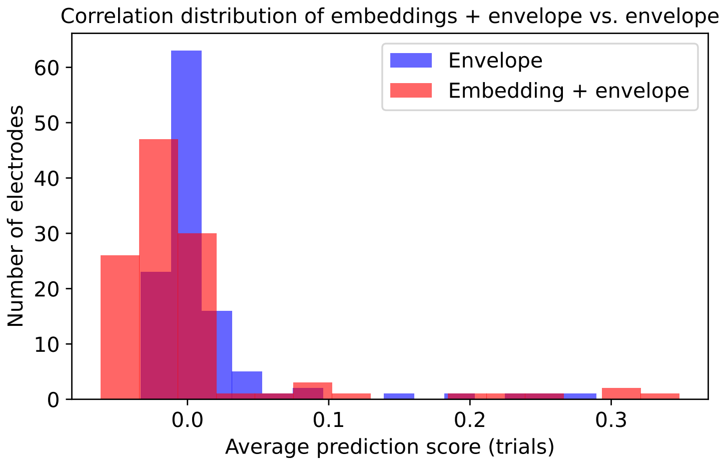


(c) Subject 3

Figure 8: **Correlation distribution of embeddings + envelope vs. envelope for the attended stream**
